# Acute physical exercise of moderate intensity improves memory consolidation in humans via BDNF and endocannabinoid signaling

**DOI:** 10.1101/211227

**Authors:** Blanca Marin Bosch, Aurélien Bringard, Maria Grazia Logrieco, Estelle Lauer, Nathalie Imobersteg, Aurélien Thomas, Guido Ferretti, Sophie Schwartz, Kinga Igloi

**Author notes:** The last two authors contributed equally to this manuscript.

## Abstract

It is well established that regular physical exercise enhances memory functions, synaptic plasticity in the hippocampus, and BDNF (Brain Derived Neurotrophic Factor) levels. Likewise, acute exercise benefits hippocampal plasticity in rodents, via increased endocannabinoids (especially anandamide, AEA) and BDNF release. Yet, whether acute exercise affects BDNF and AEA levels and influences memory performance in humans, remains to date unknown. Here we combined blood biomarkers, behavioral, and fMRI measurements to assess the impact of acute physical exercise on associative memory and underlying neurophysiological mechanisms. For each participant, memory was tested after three conditions: rest, moderate or high intensity exercise. A long-term memory retest took place 3 months later. At both test and retest, memory performance after moderate intensity exercise was increased compared to rest and high intensity exercise. We also show that memory after moderate intensity exercise benefited from exercise-induced increases in both AEA and BNDF levels: while AEA boosted hippocampal activity during memory recall, BDNF enhanced hippocampal memory representations and long-term performance. These findings confirm previous results on the benefits of acute exercise towards memory consolidation and, by including the contribution of key biomarkers, extend them by explaining neural plasticity mechanisms mediating cognitive enhancement.

**Significance statement:** Here we show that cycling for half an hour at moderate intensity after encoding new memories improves their retention when tested right after sport, and also 3 months later. This boost in memory performance occurs selectively for moderate intensity exercise, and is not observed after high intensity cycling. We report that exercise-induced memory enhancement is dovetailed by the activation of the hippocampus in the brain, and by an increase in blood concentrations of endocannabinoids (molecules involved in the feeling of euphoria after exercise) and brain-derived neurotropic factor (BDNF). Consistent with the role of the latter in neural plasticity mechanisms, we show that exercise-induced BDNF increase favors the long-term stabilization of memory representations across brain networks (i.e. 3 months after the first memory test).

## Introduction

Regular physical exercise is a lifestyle factor, which benefits neurocognitive functions and brain plasticity (1) at all ages, and may possibly reduce the risk of cognitive decline associated with Alzheimer’s disease (2). Studies in animals support the fact that voluntary regular exercise fosters neurogenesis in the adult hippocampus and improves learning and memory capacities (3). Adult neurogenesis in the human hippocampus has been repeatedly suggested (4,5), albeit being recently questioned (6). Several lines of evidence converge to suggest that enhanced hippocampal synaptic plasticity is mediated, at least in part, by brain derived neurotrophic factor (BDNF)(7). Specifically, physical exercise increases the levels of BDNF mRNA and protein in the hippocampus and other brain regions (7), and blocking BDNF action in the hippocampus hinders the beneficial effect of exercise on memory (8).

Human studies have primarily focused on the long-term effects of exercise on BDNF and cognition. Yet, measuring BDNF levels before and after a period of regular physical training does not account for the kinetics of the upregulation of this growth factor, which is thought to be predominantly fast and transient (9). In particular, BDNF levels are known to rapidly increase in hippocampal subfields in response to exercise (10), together with enhanced long-term potentiation (LTP) and synaptic plasticity (11). These effects may mediate memory enhancement on the timescale of a few hours (12). Associated with LTP induction, exercise also rapidly affects fine cell morphology, especially by increasing the number and size of hippocampal dendritic spines considered to support changes in synaptic strength (13).

In addition, physical exercise yields a rapid increase in circulating endocannabinoids, which act on cannabinoid receptors CB1 and CB2 (14-16). Work in animal models have implicated endocannabinoid signaling in exercise-induced adult hippocampal neurogenesis (17) and plasticity mechanisms (18). Endocannabinoids directly mediate different forms of retrograde plasticity (19) and can also modulate non-endocannabinoid-mediated forms of plasticity including LTP (20) and BDNF signaling (21). One recent study directly linked endocannabinoid levels to memory enhancement and hippocampus function in mice by showing that blocking CB1 receptor in the hippocampus disrupted spatial memory performance whereas artificially elevating endocannabinoid concentrations in sedentary animals increased BDNF levels and memory (22).

In humans, short periods of exercise, or acute exercise, were reported to have diverse effects on learning, memory, and cognition in humans, ranging from positive to detrimental (1,23-26). These inconsistent results may primarily be attributable to the use of different exercising intensities and/or overall poor quantification of exercise intensities (27). In a previous behavioral study, we showed that moderate intensity exercise boosted associative memory performance (26). The main aims of the present study were (i) to confirm these effects using an individually-defined calibration of moderate physical effort (corresponding here to cycling during 30 minutes at 65% of the maximal cardiac frequency measured during VO2max), and (ii) to unravel the underlying blood biomarker and neuroimaging correlates. Based on animal data (reviewed above), we hypothesized that endocannabinoids and BDNF influence hippocampal functioning after acute physical exercise in humans. As mentioned above, different exercising intensities have been used in previous work, but were rarely compared and often poorly characterized. We therefore added an exploratory high intensity exercising condition to clarify whether the beneficial effects of exercise on memory performance are specific to moderate intensity or whether they may also be observed for a high exercising intensity. During the high intensity condition participants cycled during 15 minutes at 75% of their maximal cardiac frequency, which corresponds to an effort level above the ventilatory threshold.

Here we assessed the influence of different intensities of physical exercise on memory in 18 participants, by comparing performance during three separate sessions of moderate intensity exercise, high intensity exercise, and rest period (according to a cross-over randomized within-subjects design). We used a hippocampus-dependent associative memory task (26,28) in which participants learned 8 series of 6 successive pictures. Participants first saw the 8 series once during an encoding session (Fig. 1B), followed by a 2-alternative forced choice (2AFC) learning session with feedback (Fig. 1C **– right panel**). After exercise or rest, associative memory was tested again using a 2AFC on pairs of pictures with different relational distances (direct, inference of order 1 and order 2; Fig. 1C). Sixteen control trials (used in the decoding analysis) were also included in which the depicted elements were of a given color (red, blue or green) and participants had to choose among two pictures which one was of the same color as the target picture. Blood samples were collected before and after each period of exercise or rest to measure endocannabinoids and BDNF levels. We also tested the effects of acute physical exercise on long-term memory during a surprise memory retest 3 months after the last experimental visit (Fig. 1A). Functional MRI (fMRI) data was acquired during memory encoding, learning, test, and retest, and were analyzed using SPM12 (see Methods), the results from test and retest sessions are reported here. In line with our previous results (26), we hypothesized that moderate intensity exercise would yield the largest benefits, especially at immediate test. Further, we expected that such memory benefits would be associated with exercise-related changes in BDNF and AEA levels. AEA is known to have transient effects due to its rapid degradation by metabolic enzymes (29), whereas the reported effects of BDNF are generally long-lasting (4). We therefore predicted that increases in BDNF levels may underlie long-term memory effects.

**Figure 1 –.**
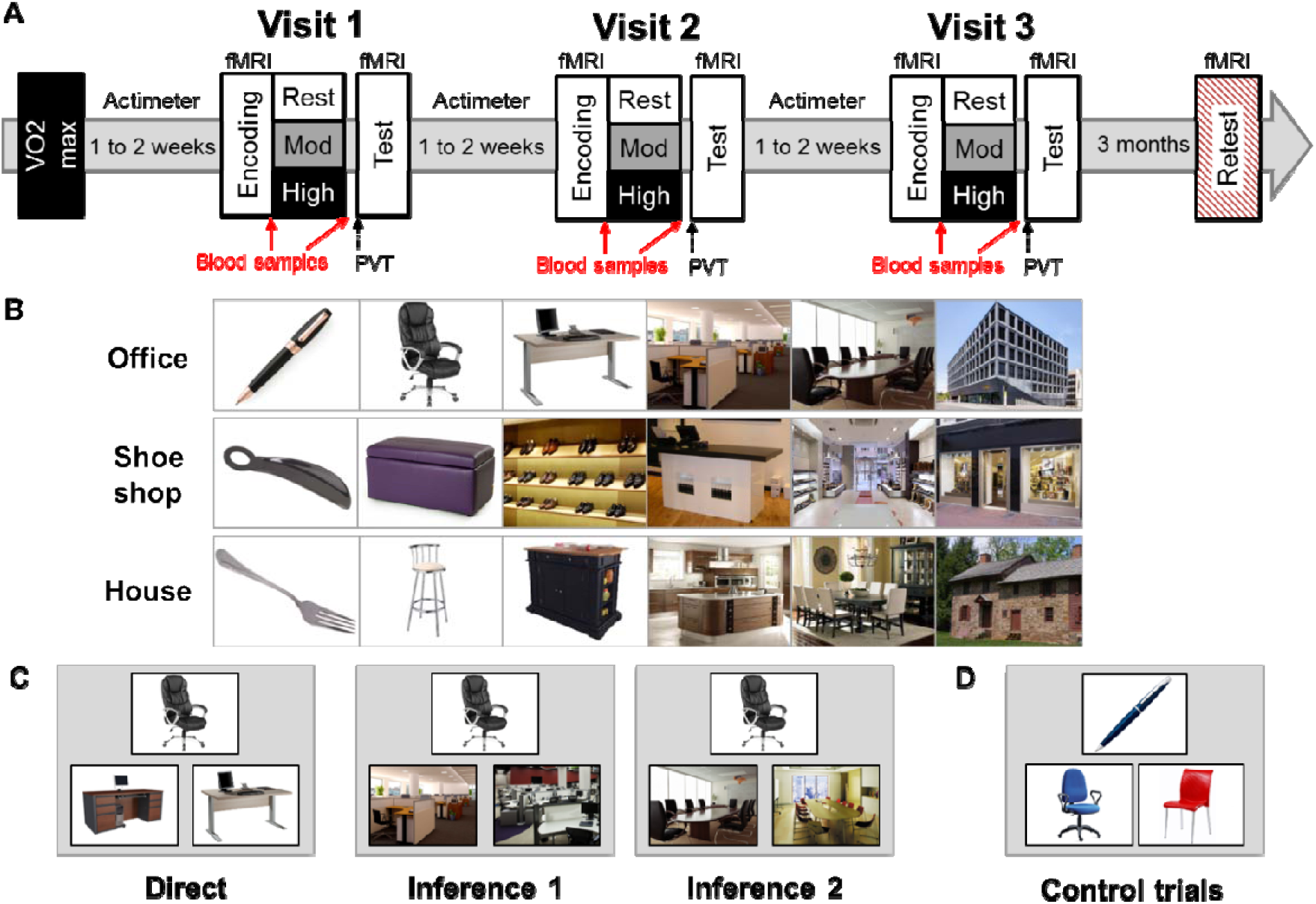
Experimental design. **A)** Overview of the experimental protocol composed of five visits: a VO2max visit, followed by three experimental visits, and a retest visit performed three months after the last experimental visit. All experimental visits started at 9AM and were composed of two MRI sessions (encoding and test) separated by a physical exercise or rest session. Physical exercise was either of moderate intensity (30 minutes cycling at 60% of FcMax) or of high intensity (15 minutes cycling at 75% of FcMax). Blood samples were collected twice in each experimental visit, before and after exercise or rest. PVT and POMS questionnaire were administered after exercise or rest. **B)** Examples of series of pictures for each theme (top row: office, middle row: shoe shop, bottom row: house). **C)** Examples of direct trials (left), inference of order 1 (middle), and 2 trials (right). Direct trials were used during the learning, test, and retest sessions, inferences 1 and 2 trials were used during test and retest sessions. **D)** Example of control trials.

## Results

### Test

To test our main prediction about the effect of physical exercise on memory and provide a replication of our previous behavioral findings (26), percentage of correct trials (% correct) and efficiency data (see Methods) from the test session were analyzed using repeated-measures ANOVAs with Exercising Condition (rest, moderate intensity exercise, high intensity exercise) as repeated measures and Relational Distance (direct, inference 1, inference 2) as within-subjects factor. We report a main effect of Exercising Condition for both % correct (F(2, 102)=4.01, p=0.02) and efficiency (F(1, 102)=8.62, p<0.01), but no effect of Relational Distance and no interaction (all p>0.05; Fig. 2A). Post-hoc analyses revealed that increased % correct and higher efficiency after the moderate compared the rest Exercising Condition (p_mod-rest_=0.02 and <0.01, respectively), while efficiency was also higher after moderate intensity exercise compared to high intensity exercise (p_mod-high_<0.01).

**Figure 2 –.**
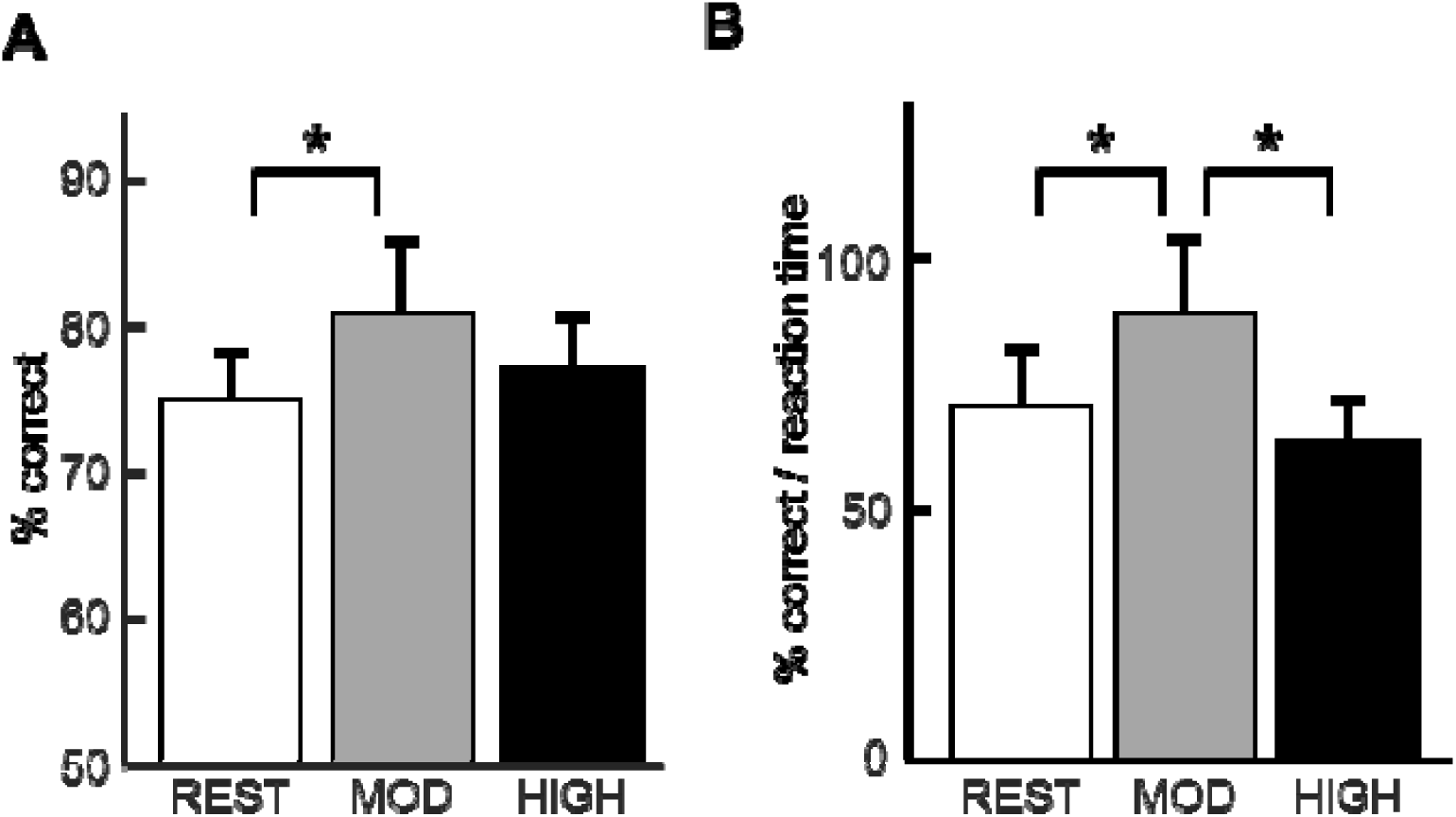
Memory performance at test. **A)** % correct: higher proportion of the percentage of correct trials after moderate intensity exercise than after rest. **B)** Efficiency (% correct / reaction time): higher efficiency after moderate intensity exercise than after rest and high intensity exercise. All bar plots represent mean +/- SEM.

### Blood samples

We measured changes in endocannabinoids and BDNF levels from the blood samples collected right before and after the rest and exercise sessions (see *SI Appendix* for details). Repeated-measures ANOVAs were performed for each biomarker with Exercising Condition (rest, moderate, high) as a within-subjects factor. For AEA, a main effect of Exercising Condition (F(2, 34)=39.25, p<0.01; Fig. 3A) was found. Post-hoc analyses revealed that AEA levels were lower after rest than after physical exercise (p_rest-mod_<0.01, p_rest-high_<0.01), with no difference in AEA levels after moderate and high intensity exercise (p_mod-high_>0.05). Please note that AEA during the rest condition decreased from the first (baseline) to the second (post-rest) measurement, hence resulting in a negative differential value. This decrease is consistent with known circadian fluctuations in AEA, whereby AEA levels increase during sleep and decrease throughout the day (30). For the endocannabinoid 2-arachidonoylglycerol (2-AG), there was no effect of Exercising Condition (F(2, 34)=2.90, p>0.05), consistent with previous descriptions in the literature (15). For BDNF, we report a main effect of Exercising Condition (F(2,34)=4.78, p=0.01; Fig. 3D). Post-hoc analyses revealed that, BDNF levels after moderate and high intensity exercise differed from after rest (p_rest-mod_=0.045, p_rest-high_=0.01).

**Figure 3 –.**
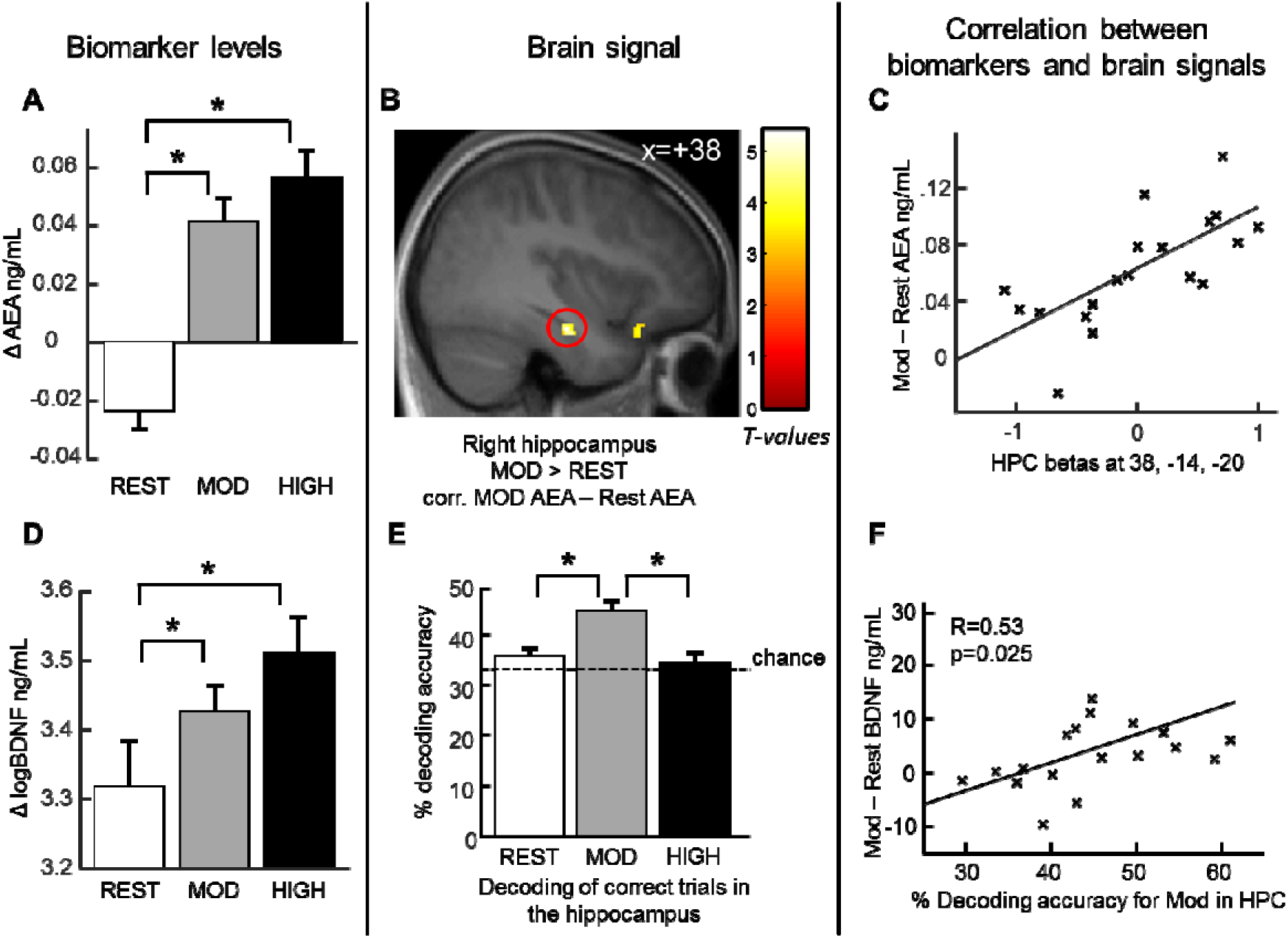
Increased biomarker levels correlate with hippocampal brain signals after moderate intensity exercise. **A)** Increased Anandamide level (AEA) after moderate and high physical exercise compared to rest. For all Exercising Conditions Δ AEA corresponds to the difference in AEA between the second blood sample (after exercise or rest) and first blood sample (before exercise or rest). **B)** Increased right hippocampal response [z-score=3.72 (38, −14, −20), p<0.05 SVC] for correct responses after moderate exercise compared to correct responses after rest correlated with the increase in AEA level after moderate exercise. **C)** Correlation between the hippocampal beta values and AEA. **D)** Increased BDNF levels after moderate and high intensity exercise compared to after rest. For all Exercising Conditions Δ BDNF corresponds to the difference in BDNF between the second blood sample and the first blood sample. For display purposes, we represent ΔlogBDNF. **E)** Higher sensitivity in decoding accuracy of correct trials in the bilateral hippocampus after moderate exercise than rest and high intensity exercise. **F)** Positive correlation between decoding accuracy in the hippocampus and increase in BDNF level after moderate intensity exercise. Activation map displayed on the mean T1 anatomical scan of the whole population. For display purposes, hippocampal activations are thresholded at p<0.005.

### Psychomotor vigilance test (PVT) and Profile of Mood States questionnaire (POMS)

We administered the PVT and POMS just before the MRI test session (i.e. about 45 min after rest or exercise) to monitor possible condition-dependent differences in vigilance and mood at the time of the test session. For PVT, we replicate our previous results (26) showing no difference in PVT as a function of Exercising Condition (rest, moderate, high), neither in mean or median reaction times, number of lapses, or number of false alarms (one way repeated measures ANOVAs, all p>0.05). For POMS, we report no difference for any of the measured categories (fatigue, tension, confusion, vigor) as a function of Exercising Condition (all p>0.05), suggesting that the physical exercise sessions did not result in significant lasting mood or vigilance changes.

### Functional MRI results

#### Test

We first conducted a standard general linear model analysis with the data collected during the memory test after rest, moderate intensity exercise, and high intensity exercise modelled as separate sessions. Within each session, we considered correct trials according to Relational Distance (direct, inference 1, inference 2) and control trials as four separate regressors of interest, and included incorrect trials as an additional regressor (Fig. 1). When comparing high Relational Distance to low Relational Distance (inference 2 > direct trials) across all sessions, we found increased activity in the right hippocampus [z score=4.35 (18, −38, −8), p<0.05 SVC, see Methods], bilateral parahippocampal gyrus and precuneus (see **Fig. S1** and **Table S1** for exhaustive list of activated regions). No region was activated (at a threshold of 0.001 unc.) when comparing inference 1 to direct trials, and inference 2 to inference 1 trials. Comparisons between Exercising Conditions and interactions between Relation Distance and Exercising Conditions did not yield any significant activation either.

As it is known that AEA has a rapid effect on synaptic plasticity in the hippocampus, we tested whether the observed difference in AEA across Exercising Conditions might exert a modulatory influence on brain activity. We thus added individual AEA change as a cofactor in the second-level analyses comparing Exercising Conditions. We found that the increase in AEA after moderate intensity exercise (vs. rest) correlated with the activation in the right hippocampus [z-score=3.72 (38, −14, −20), p<0.05 SVC] (Fig. 3B). A similar modulation of hippocampal activity was found for high intensity exercise (vs. Rest) [z-score=4.08 (32, −24, −18), p<0.05 SVC], suggesting that AEA increase correlated robustly with hippocampal activation (**Fig. S2**).

Next, we used a decoding approach to test whether exercise would affect the coherence of the fine-grained neural representation of correct, incorrect and control trials within the bilateral hippocampus. To test for this, we applied a similar procedure as Van Dongen et al ((23); see *SI Appendix* and Methods section), to classify each single trial, from voxelwise hippocampal activity, into one of three possible outcomes (correct, incorrect or control trial), with a chance level at 33.33%. Focusing on correct trials, we report that decoding accuracy was above chance level after moderate intensity exercise, but at chance level after rest and high intensity exercise (Fig. 3E). Post-hoc analyses further showed that decoding after moderate intensity exercise was higher than after both rest and high intensity exercise (p_mod-rest_<0.001, p_mod-high_ <0.001, depicted on Fig. 3E). We obtained similar results when we performed the classification on activity from the left or the right hippocampus separately (see **Fig. S3**).

Because BDNF is known to specifically enhance plasticity mechanisms in the hippocampus, we tested whether BDNF levels may affect the neural representation as measured with decoding results. We report a positive correlation between BDNF enhancement during moderate intensity exercise (calculated as the difference between moderate and rest BDNF values, with baseline values subtracted for each visit) with decoding accuracy after moderate intensity exercise (R=0.53, p=0.02), but not high intensity exercise (p>0.05); Fig. 3F.

### Retest

All participants came back for a long-term memory retest session three months later. A repeated-measure ANOVA was performed with Exercising Condition (rest, moderate, high) as within-subjects factor, that revealed a main effect of Exercising Condition (F(2, 34)=3.32, p=0.048). Post-hoc analyses showed that this main effect was due to participants performing better for the associations learned during the moderate exercise session as compared to those learned during rest session (p_mod-_ rest=0.04); Fig. 4A. Moreover, only the trials from the moderate exercise session were remembered above chance level three months later (t(17)=2.31, p=0.03).

**Figure 4 –.**
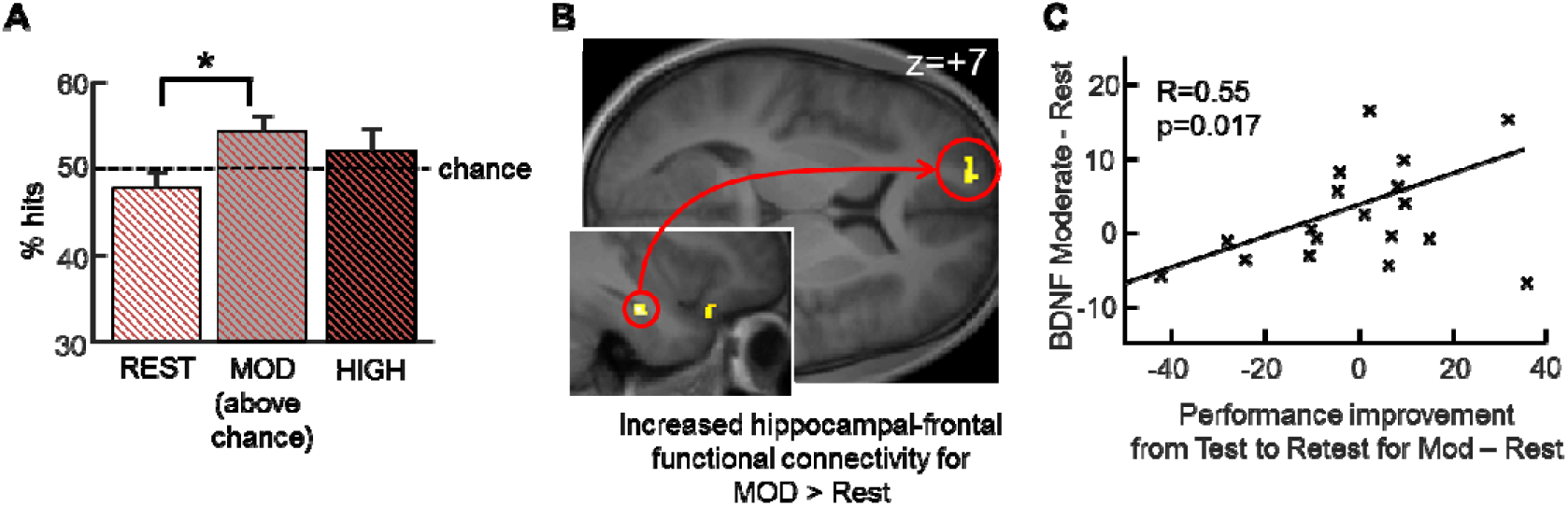
Better long-term memory for associations learned after moderate physical exercise, related to prefrontal activation and BDNF signaling. **A)** Better performance for pictures learned during the moderate intensity visit than for pictures learned during the resting visit. Performance after moderate exercise is significantly above chance level. **B)** PPI for the retest session, using the seed in the left hippocampus from Figure 3B. Increased functional coupling with the left superior frontal gyrus [z-score=3.16 (−16, 62, 6), p<0.001], selectively after moderate exercise compared to after rest. **C)** Performance improvement from test to retest for moderate exercise compared to Rest correlates with BDNF enhancement from moderate exercise to rest.

We then asked whether physical exercise had some long-term effects on the functional coupling between the hippocampus and other brain regions during the processing of associative memories. We therefore performed a psychophysiological interaction analysis (see Methods) taking as seed region the right hippocampal region that initially showed increased activity for moderate intensity vs. rest as a function of AEA levels (38, −14, −20; see Test results above). We observed increased functional connectivity between the hippocampal seed region and the left superior frontal gyrus [z-score=3.10 (−16, 62, 6), p<0.001 unc.] when participants were exposed to associations learned during the moderate intensity exercise session (compared to rest; Fig. 4B).

Based on the existing evidence that BDNF contributes to neurogenesis and synaptic plasticity (4, 7, 8), we also hypothesized that changes in BDNF levels during the moderate intensity exercise condition (i.e. condition associated with the highest benefit for short-term memory consolidation and hippocampal representations) may potentially promote long-term memory retention. This hypothesis was tested by correlating individual changes in BDNF levels (for moderate intensity vs. rest) to delayed performance increase (i.e., from test to retest) for items initially learned during the moderate vs. rest condition. We report a significant positive correlation (R=0.55, p=0.02; Fig. 4C), while the same correlation for high intensity exercise was not significant (R=0.31, p=0.20). These results suggest that BDNF increase after moderate intensity exercise may contribute to durable memory enhancement.

## Discussion

We show here that one session of moderate intensity physical exercise compared to a period of rest enhanced associative memory, both at immediate test (2 hours after encoding) and at long-term retest (three months later). These effects may be mediated by the endocannabinoid AEA and the growth-factor BDNF, whose respective concentrations increased after acute exercise. Accordingly, during the short-term test, the increase in plasma AEA concentration correlated with hippocampal activity when associative memories were recalled, and BDNF increase correlated with decoding measures within the hippocampus. Moreover, BDNF increase during moderate intensity physical exercise correlated with better performance at long-term retest. We did not observe such memory benefits for a session of high intensity physical exercise, i.e. when effort levels surpassed the ventilatory threshold. Overall, we show that acute physical exercise at moderate intensity has long-lasting positive effects on the consolidation of associative memories in healthy young human adults. Below, we discuss the neurophysiological mechanisms that could explain these important findings.

### Biomarker mechanisms underlying the effects of acute exercise on hippocampal plasticity

In a recent study in rodents, Fuss et al. (31) demonstrated that physical exercise induces an acute increase of AEA measured in the plasma, with direct effects on CB1 receptors in the brain. Note that in the same study cerebro-spinal fluid measures did not capture increases in AEA, consistent with AEA being very rapidly metabolized in the brain (29). These results support the fact that plasma measures of AEA, as we performed here, offer a reliable index of AEA activity in the central nervous system. Another rodent study directly linked endocannabinoid signaling to hippocampal memory function, by showing that selectively blocking CB1 receptors in the hippocampus abolished exercise-induced memory effects (22). This study also demonstrated that artificially increasing AEA concentrations (by blocking the Fatty Acid Amine Hydrolase, the enzyme responsible for breaking down AEA) in the hippocampus of sedentary mice mimicked the effects of physical exercise and increased memory performance. Together, these rodent studies elucidate the neurophysiological mechanisms underlying our novel finding that AEA increase in human plasma may reflect direct effects of physical exercise on brain activity, especially in the hippocampus.

Traditionally, BDNF has been linked to effects of regular physical exercise, although it is known that BDNF gene expression is upregulated both after acute and after chronic physical exercise in rodents (32). It is widely acknowledged that BDNF enhances synaptic plasticity, especially via LTP (12), which can be induced in a few minutes and critically contributes to memory consolidation (33).Here we show that the effects of one single session of exercise may differentially affect both short and long-term memory retention. On the one hand, BDNF increase after acute moderate physical exercise correlated positively with decoding accuracy of memory items in both hippocampi immediately after exercise (test session). On the other hand, BDNF increase (at test) also correlated with long-term memory differences (at retest) due to the initial exercising conditions. Specifically, those participants who exhibited larger increases in BDNF levels at test after the moderate intensity Exercising Condition remembered the learned associations better at retest three months later.

### Acute moderate but not high intensity exercise benefits memory consolidation

While characterizing the impact of exercise intensity on cognitive functions is critical for health recommendations, dementia prevention programs and rehabilitation strategies, the reported effects remain inconsistent. Some studies suggest that high intensity training is most efficient (23,25) while other studies, especially meta-analyses, indicate that moderate intensity exercise might have more impact (34), and a recent report show that very mild intensity exercise (at 30% of maximal cardiac frequency (FcMax)) may already benefit hippocampal memory function (24). Here we aimed at clarifying this important issue by using a cross-over randomized within-subjects design according to which each participant was tested at a moderate and at a high intensity (plus a resting, baseline condition) across distinct sessions where associative memory was also tested. Importantly, here we determined moderate and high intensity exercise levels with reference to each participant’s individual ventilatory threshold. This threshold was measured using a VO2max procedure (see Methods), which is a gold-standard in human physiology research (see 35 for review). Moderate intensity corresponded to exercise below the ventilatory threshold (65% of individual VO2max) and high intensity corresponded to exercise above the ventilatory threshold (75-80% of individual VO2max). Here we found strong beneficial effects of moderate intensity exercise on memory, with a clear difference compared to rest, while the effects of high intensity exercise appeared to be more complex. Some evidence suggests that while aerobic exercise training (i.e. below ventilatory threshold) is beneficial for hippocampal functioning, high intensity training is not (36). One plausible explanation is that acute high intensity exercise may induce a physiological stress response (for example a strong increase in cortisol levels) which can impair memory for previously learned stimuli (37,38). In this article, rather than concentrating on the clear-cut positive results between moderate intensity exercise and rest, we decided to present the findings from both intensities and carefully discuss below the possible reasons for the differential effects of moderate and high intensity exercise. We hope that our results and the ideas raised in our discussion will fuel future debates and investigations in the scientific community.

Here we observed that moderate levels of exercise intensity increased both BDNF and AEA levels and optimized cognitive processes. By contrast, although high intensity physical exercise further increased the measured concentrations of BDNF and AEA, performance did not follow this increase. This observation suggests that large increases in BDNF and AEA concentrations might not be as beneficial for memory performance. Please note that while, at this point, we cannot exclude that there is no link between BDNF and AEA levels and memory consolidation, this latter hypothesis is not prevalent in the current literature. In line with our findings, Mamounas et al.(39) showed that the BDNF dose-response curve follows an inverted U-shape with intermediate concentrations of BDNF yielding best results for sprouting of serotoninergic neurons in the rodent hippocampus. For AEA, one study using exogenous AEA administration suggested that related anxiolytic effects also follow an inverted U-shape dose-response curve with highest concentrations (measured in the periaqueductal gray) being less effective (40). The findings of the present study also suggest that submaximal concentrations of both molecules (as obtained after moderate intensity exercise) yield best effects on neurocognitive functions, here for hippocampal-dependent memory formation. As mentioned above, we cannot exclude that other biomarkers may also contribute to the observed effects, such as for example a large increase in cortisol after high intensity exercise, which may be detrimental for memory consolidation (37,38).

### Long-term consequences of acute physical exercise on memory consolidation

Lasting effects of physical exercise are established for regular physical exercise protocols, involving several months of training (4,41). Most of these studies focused on the possible protective effects of physical exercise in ageing and dementia. Yet, acute physical exercise has also been reported to have positive short term cognitive effects (23,26), albeit not always found in tasks involving hippocampus-dependent memory (42). Long-term effects of acute physical exercise (at the scale of several months as we tested here) have to our knowledge not been investigated in humans so far. Here we found long-lasting effects on memory retention selectively for moderate intensity exercise (i.e. below the ventilatory threshold), which were dovetailed by increased connectivity between hippocampus and prefrontal cortex.

## Conclusion

We show that acute moderate intensity physical exercise significantly increased associative memory performance both at short and long term. At short term, hippocampal activation correlated with endocannabinoid AEA while enhanced hippocampal memory representations were associated with a modulation of BDNF. At long term, three months after encoding, memory effects were related to BDNF increase induced by moderate intensity exercise. High intensity exercise did not have such beneficial effects. We conclude that a single session of moderate physical exercise boosts associative memory formation.

## Methods

### Participants

We included 20 healthy young male volunteers in this study. All participants gave written informed consent and received financial compensation for their participation, which was approved by the Ethics Committee of the Geneva University Hospitals. Two participants had to be excluded from all the analysis for non-compliance with experimental requirements. The remaining 18 participants were between 18 and 34 years old (mean age +/- standard error: 23.03+/-0.92 years). All participants were right-handed, non-smokers, free from psychiatric and neurological history, and had a normal or corrected-to-normal vision. Please note that, for the present experimental design, we estimated the required sample size based on the results from our previous study (26) (see *SI Appendix* for details).

### Experimental procedure

Participants first came to the lab for a VO2max procedure. During this visit, participants also performed a habituation session of the associative task. Those participants with a VO2max within the required ranges (see *SI Appendix*) were invited to come back for three experimental visits separated by one to two weeks, according to a within-subjects design with the three Exercising Conditions (rest, moderate intensity exercise, high intensity exercise) randomly counterbalanced across participants.

For each visit, participants arrived at 08:00 AM on an empty stomach, and had a controlled breakfast with an experimenter (*SI Appendix*). At 09:00 AM, participants were comfortably installed in the scanner, and started the encoding part of the associative memory task (see below and Fig. 1A) while fMRI data was acquired. At 09:50 AM a qualified medical doctor took a first blood sample. At 10:00 AM participants were equipped with a Polar RS800CX N device to measure heart rate and asked to rest or exercise. For the two exercise conditions, participants pedaled on a cycle ergometer (Ergoline GmbH, Bitz, Germany), the pedaling frequency was kept between 60 and 80 cycles per minute, which was shown on a small screen in front of the participant. For moderate intensity exercise, each participant pedaled for 30 minutes, with the load of the ergometer set so that the cardiac frequency of the participant would be at 60% of his FcMax. For high intensity, participants first warmed up for 2 minutes at 50% of FcMax then the load was progressively increased over 1 minute to reach 75% of FcMax. Participants pedaled at this intensity for 15 minutes then they pedaled again at 50% of FcMax for 3 minutes to cool down. For both exercise conditions, the experimenters checked cardiac frequency every 3-5 minutes to adjust the resistance of the ergometer if necessary. For the rest condition, participants sat on a chair and were allowed to quietly look at magazines for 30 minutes. To minimize interference with memory, we carefully selected these magazines so that they were mainly composed of pictures, and that there was little to be learned from their content. We purposefully did not let participants watch a movie during rest to minimize motor imagery. At 10:30 AM, the medical doctor took a second blood sample and fifteen minutes after this, participants performed a Psychomotor Vigilance Task (PVT) followed by the Profile of Mood States (POMS) questionnaire. These latter two measures were acquired as control measures to exclude that difference in fatigue and mood states may explain our memory findings so they were administered when heart rate and other physiological conditions were back to baseline, close in time to the second fMRI session when memory was tested.

At 11:30 AM, participants underwent a second fMRI session during which memory for the associative task was tested. A surprise retest fMRI session took place three months later where participant’s memory was tested again; no blood samples were taken at this time point.

#### Associative memory task in fMRI

We adapted an associative memory task (26,28) consisting of two parts: encoding and test, separated by an exercise (moderate or high intensity) or rest period (Fig. 1A). To avoid interference across experimental visits for this within-subjects design, we showed different pictures belonging to three specific themes at each visit: “office”, “shoe shop” or “house” (one theme per visit). The pictures in each theme for the experimental visits were matched in difficulty and counterbalanced across Exercising Conditions and visits (Fig. 1B). Note that for the habituation session of the task, participants had to memorize 5 series of a “swimming pool” theme.

During the encoding session, participants were first shown 8 series of 6 pictures passively once. Then, participants were presented with the first picture of the series alone (e.g., pen, for the “office” theme; Fig. 1B). Then, the same first picture was presented in the upper half of the screen together with two options for the second picture in the series (chair) in the lower half of the screen, one being the correct next picture and the other picture being from a different series (as depicted on the left panel of Fig. 1C). Participants had to select the correct next picture by pressing a left or right button. The correct picture was then shown (providing a feedback for each trial), followed by this same picture together with the two next options for the third picture in the series (desk). This continued until the last picture in the series (office building) (*SI Appendix* for details).

During the test session, participants were presented with one cue picture and two other pictures, among which they had to select the one belonging to the same series as the cue picture. The two options could represent the immediate next item in the series (direct trials) or could be separated by one or two items from the cue picture (inference of order 1 or order 2 trials; Fig. 1C). All types of trials were shown in a randomized order, and were presented in the same format and with the same timeframe as during learning, except that no feedback was provided. In this session, 16 trials of the control “color” task were also included.

For the delayed retest session, all 18 participants came back for a surprise retest in fMRI three months after the last experimental visit. Participants did not know at test that there would be a retest session. The task was identical to the test sessions, except that pictures of all three themes were now mixed in a random order. For time constraints, we included for each of the eight sequences of pictures of all three themes (24 sequences) two direct trials, two inference order 1 trials and one inference order 2 trial (5 trials) totaling 120 trials overall.

### Functional MRI data acquisition and analysis

MRI data were acquired on a 3 Tesla MRI scanner (SIEMENS Trio® System, Siemens, Erlangen, Germany) with a 32-channel head coil. T2*-weighted fMRI 2D images were obtained with a multiband gradient echo-planar sequence acquiring 3 slices at a time using axial slice orientation (66 slices; voxel size, 2 x 2 x 2mm; repetition time (TR) = 1880ms; echo time (TE) = 34ms; flip angle (FA) = 60°). A whole-brain structural image was acquired at the end of the first test part with a T1-weighted 3D sequence (192 contiguous sagittal slices; voxel size, 1.0 x 1.0 x 1.0mm; TR = 1900ms; TE = 2.27ms; FA = 9°). Continuous measures of heart rate and breathing rhythm were acquired using a Biopac (Biopac Systems, CA93117, USA).

#### Conventional fMRI analysis

Functional images were analyzed using SPM12 (Wellcome Department of Imaging Neuroscience, London, UK). This analysis included standard preprocessing procedures (*SI Appendix*). We performed corrections to regress out potential physiological artifacts from heart rate and breathing using Retroicor (43) and RVHcorr (44), respectively. A general linear model (GLM) approach was then used to compare conditions of interest at the individual level, each individual GLM included correct trials separated according to Relational Distance (Direct, Inference 1, Inference 2 trials), control trials and incorrect trails (pooled across Relational Distance), plus 6 movement regressors, 5 heart rate regressors and 1 breathing regressor as regressors of non-interest. Then, contrasts between conditions of interest from each participant were entered a second-level random-effects analysis. All activations are reported at p<0.001 with a cluster size of 10 voxels and relevant regions, especially the hippocampus, survived small-volume correction (SVC) for familywise error (p < 0.05) using volumes based on the Anatomy toolbox of SPM12 (SPM Anatomy toolbox 2.2, Forschungszentrum Jülich GmbH). Coordinates of brain regions are reported in MNI space. See *SI Appendix* for PPI and decoding analyses.

## Acknowledgements

We are grateful to the Brain and Behaviour Laboratory of the University of Geneva (Geneva, Switzerland) for providing help and scanning facilities. We thank Emrah Duezel for useful comments on this manuscript.

## Author contributions

B.M.B, A.B, G.F., S.S. and K.I. designed research; B.M.B, A.B., M.G.L., N.I. and K.I performed research; B.M.B., A.B., M.G.L.,E.L., A.T., S.S. and K.I. analyzed data; and all the authors wrote the paper.

