## Supplemental Information for "Acute physical exercise of moderate intensity improves memory consolidation in humans via BDNF and endocannabinoid signaling"

### SI Appendix

### SI results

#### Learning

The percentage of correct responses (% correct) and efficiency (i.e. % correct divided by reaction time) were analyzed using repeated measure ANOVAs with Learning Blocks (block 1, block 2, block 3) and Visit theme (office, shoe shop, house) as within-subjects factors. Both analyses revealed a main effect of Block (hit rate: F(2, 34)=23.16, p<0.001; efficiency: F(2, 34)=17.64, p<0.001), consistent with a progressive learning of the associations, but no effect of Visit theme (hit rate: F(2, 34)=1.92, p=0.16; efficiency: F(2, 34)=0.16, p=0.85) and no interaction (hit rate: F(4, 68)=0.56, p=0.69; efficiency: F(4, 68)=0.41, p=0.80). Importantly, there was no main effect and no interaction with subsequent physical exercise neither for hit rate nor for efficiency (all p>0.05) when this factor was added as repeated measure to the previous ANOVAs. Overall, during the third block, participants reached a high level of performance (hit rate ± standard error: 86.94 ± 1.63%), suggesting a good encoding of the series, well above chance level.

#### Test

In addition to the reported results in the main text, we tested for possible effects of Visit Theme. We therefore performed an additional repeated-measures ANOVA with Visit Theme (office, shoe shop, house) and Relational Distance (direct, inference 1, inference 2) as within-subject factors. We report no effect of Visit Theme (F(2, 102)=1.40, p=0.25) and no effect of Relational Distance and no interaction effect (both p>0.05).

Blood samples: The first blood sample served as a baseline measure. This is especially important for endocannabinoid measures, which are known to substantially fluctuate according to diet and other environmental factors. The values reported in the main text were obtained for each visit by subtracting the first blood sample (baseline measure) from the second blood sample.

Psychomotor vigilance test (PVT) and Profile of Mood States questionnaire (POMS): Both PVT and POMS were performed just before the MRI test session. This was more than half-an-hour after the end of the exercising/rest period when heart rate and other physiological measures were back to baseline. The PVT and POMS were used to monitor possible condition-dependent differences in vigilance and mood at the time of the test session (1). For PVT, we replicate our previous results (2) showing no difference in PVT as a function of Exercising Condition (rest, moderate, high), neither in mean or median reaction times, number of lapses, or number of false alarms (one way repeated-measures ANOVAs, all p>0.05). This suggested that participants did not significantly differ in vigilance state 45 minutes after rest or physical exercise. Concerning the POMS questionnaire, we used a short version composed of 38 questions assessing tension, fatigue, vigor, confusion with 7 to 9 questions for each category and including an additional 7 dummy questions. Each question has 5 levels: 0=not at all, 1= a little, 2=moderately, 3=quite a lot, 4=extremely. The POMS score for each category is the sum of the scores for the corresponding questions. For POMS, we report no difference for any of the measured categories (fatigue, tension, confusion, vigor) as a function of Exercising Condition (all p>0.05), suggesting that the physical exercise sessions did not result in lasting mood changes.

Correlation with general fitness level: To verify that individual fitness levels did not significantly affect our main results, we tested whether behavioral performance (i.e. % correct and efficiency) and biomarker levels (i.e. AEA, BDNF), correlated with VO2max measures, but found no correlation (all p>0.05).

Heart rate and breathing analysis: Participants’ heart rate measured by a Polar cardiofrequencemeter (Polar RS 800 CX, Polar, Finland) during the rest period was at 33.4+/-4.0% of their maximal heart rate as assessed by the VO2max procedure (see Methods). During moderate and high intensity physical exercise, participants pedaled at 68.7+/-1.1 % and 77.7+/-1.9% of their maximal heart rate, respectively. A one way repeated-measures ANOVA revealed a significant difference in heart rate between the three Exercising Conditions (rest, moderate, high intensity exercise; F(2, 34)=1672.9, p<0.001). Post-hoc analyses confirmed that the three Exercising Conditions differed from each other (all p<0.001). We also recorded heart rate during the test part in fMRI using the Biopac and found no difference in heart rate as a function of the Exercising Condition (all p>0.05), suggesting that all participants’ heart rate was back to baseline at test (i.e., at least 45 min after the completion of the exercise session).

Decoding approach: We performed an ANOVA on classification accuracy with Trial Type (correct, incorrect, control) and Exercising Condition (rest, moderate, high) as factors. We observed an effect of Trial Type (F(2, 34)=33.77, p<0.001) and an interaction between Exercising Condition and Trial Type (F(4, 68)=2.58, p=0.04). Separated post-hoc analyses for each trial type showed that correct trials were better classified in the moderate condition (F(2, 34)=9.43, p<0.001), while there was no effect of Exercising Condition for the control trials and incorrect trials (p=0.12 and p=0.78 respectively).

#### Retest

Retest was similar to the test sessions but it comprised five trials for each of the eight sequences of pictures of all three themes (24 sequences) amounting to 120 trials. For each sequence, two direct trials, two inference order 1 trials and one inference order 2 trials were included to equalize difficulty from all visits. For the analysis of the retest session, trials were not separated according to Relational Distance as no behavioral effect related to this was previously found.

### SI Discussion

#### Acute effect of BDNF on memory

How can we explain that an acute modulation of BDNF levels affects memory? BDNF facilitates LTP by activating signaling pathways (including MAPK and Akt) (3), promoting cytoskeleton changes (4), and enhancing protein synthesis required for vesicle trafficking and the release of neurotransmitters (5). Several studies have now confirmed that physical exercise, both per se and via BDNF signaling, boosts LTP (6, 7) via glutamatergic NMDA receptor activation. Indeed, on the one hand, physical exercise increases the expression of both NR2A and NR2B subtypes of the NMDA receptor in the hippocampus (8, 9) while, on the other hand, BDNF modulates the activity of NMDA receptors at hippocampal synapses (10). Studies in rodents have repeatedly shown that increasing NMDA-receptor-mediated plasticity is crucial for associative memory acquisition and consolidation (11, 12). In our experimental design, inducing LTP in the hippocampus (through exercise) after the encoding of new associations likely strengthens memory representations and consolidation, hence affecting pattern completion in the hippocampus for example (13), hence supporting better classification results in the exercising condition that promoted LTP (i.e. moderate exercise).

#### Possible confounding factors due to fatigue or carry-over effects of exercise

Fatigue and reduced vigilance are known to affect cognitive performance. We sought to minimize any potential effect of exercise-related fatigue (i) by scheduling the test part of the protocol 1 hour after the end of the physical exercise session and performing all experimental visits at the same time of the day (always in the morning from 8AM to 12AM); (ii) by including only participants who were exercising regularly and whose VO2max levels were above 40ml/kg/min, so that exercise intensity and duration would not be exhausting for them. We also specifically measured fatigue and vigilance levels in our participants and found that neither POMS scores for fatigue nor PVT did differ after moderate or high intensity exercise. We also checked that heart rate and breathing rhythm of all our participants were back to baseline levels when the test session started. Finally, to exclude any contaminations of heart rate or breathing on our fMRI data, we carefully regressed out these effects using Retroicor (14) and RVHcorr (15, 16).

### SI Methods

#### Sample size

For the present experimental design, we estimated the required sample size based on the results from our previous study (2). The latter was performed on an independent sample of participants and using the same associative memory task with a similar within-subject design, where we found a significant behavioral effect of moderate intensity exercise on memory performance (14 participants, F(1,13)=17.27, p=0.001). Using the effect size from that previous study (Cohen’s d=0.53) with an alpha level at 0.05 and power (1-beta) of 0.80, we derived an overall sample size of 15 for a two-tailed dependent-sample t-test. The present study reports results from a final sample size of 18 participants.

#### Participants

All twenty participants (initially included in the study) were within the normal ranges on self-assessed questionnaires for depression (BDI(17)), anxiety (STAI(18)), circadian typology (PSQI(19), and reported exercising regularly (at least twice per week). We only included participants whose VO2max was above 40ml/kg/min and below 65ml/kg/min (see **SI Fig4** for the distribution of VO2max measures). The lower bound was used to ensure that participants would tolerate the high intensity condition. The upper bound was needed for a homogeneous high intensity exercise condition above the ventilatory threshold (which corresponds in a cycling paradigm to about 70% of maximal cardiac frequency for moderately fit participants). In very highly trained participants, whose VO2max is above 65ml/kg/min, the ventilatory threshold is often higher, at about 90% of maximal cardiac frequency (or 80% of VO2max). Thus, an intensity defined as 75% of VO2max would not correspond to comparable difficulty levels for moderately fit vs. highly trained participants.

#### Experimental procedure

Participants were asked to keep a regular exercising schedule during at least 5 days before each visit. Moreover, they were requested to refrain from intense physical activity for the 48h preceding the experimental visits. Compliance was documented by fitness tracker (Fitbit Charge HR, Fitbit, San Francisco, USA).

For each visit, participants arrived at 08:00 AM on an empty stomach, and had a controlled breakfast consisting of coffee or tea, orange or apple juice, bread, and jam. Participants were allowed to eat as much as they desired but we controlled that they ate approximately similar amounts for all visits, they were allowed one caffeinated drink only. We did not allow them to eat any lipids to minimize inter-subject variability in endocannabinoid measures which heavily depend on lipid consumption.

Associative memory task in fMRI: During the encoding session, participants were first shown 8 series of 6 pictures one picture at a time (2000ms per picture), and were asked to encode each series as a whole (**Fig1B**). Then, they were trained on the 8 series 3 times, i.e. during three successive learning blocks. For each series, participants were shown the first picture of the series alone (e.g., pen, for the “office” theme; **Fig1B**) presented during 2000ms. Then, the same first picture was presented in the upper half of the screen together with two options for the second picture in the series (chair) in the lower half of the screen, one being the correct next picture and the other picture being from a different series (as depicted on the left panel of **Fig1C**). Participants could not answer for 2000ms, then the sentence “choose the next element” appeared on the screen and participants were instructed to press a button to give their answer when they had made their decision. Participants had to select the correct next picture by pressing the left or right button. The correct picture was then shown (providing a feedback for each trial), followed by this same picture together with the two next options for the third picture in the series (desk). This continued until the last picture in the series (office building). Additionally, two control series occurred pseudo-randomly during each block during which participants were shown a picture of a given color (red, green or blue) and had then to choose the picture of the same color (**Fig1D**). During each learning block, all 8 associative memory series and 2 control series were shown once. Stimuli were delivered and responses recorded using a MATLAB Toolbox (Cogent 2000, <http://www.vislab.ucl.ac.uk/cogent_2000.php>).

Behavioral analysis: We measured % correct and reaction times for each trial and derived efficiency as % correct divided by reaction time. % correct and efficiency were the main outcomes described in the results section. All behavioral analyses were performed using Statistica (Version 12, www.statsoft.com, StatSoft, Inc. TULSA, OK, USA). Repeated-measures ANOVAs were performed and Neuman-Keuls post-hoc comparison methods were used. Correlations were performed using the Spearman’s Rho.

VO2max measure: A maximal incremental test was performed during a preliminary visit to the laboratory, using an electrically braked cycle ergometer (Ergometrics er800S, Ergoline, Jaeger, Germany). Respiratory gas flows and ventilation were continuously measured at the mouth on a breath-by-breath basis, using a metabolic unit (K4b^2^, Cosmed, Italy), consisting of a Zirconium Oxygen analyzer, an infrared CO2 meter and a turbine flowmeter. As recommended by the manufacturer, the gas analyzers were calibrated with ambient air and with a mixture of known gases (O_2_ 16 %, CO_2_ 5 %, N_2_ as balance), and the turbine by means of a 3-l syringe. Beat-by-beat heart rate (HR) was continuously monitored by cardiotachography (Polar RS 800 CX, Polar, Finland). Gas exchange variables (V˙O2, V˙CO2, V˙E, and RER) were continuously recorded on a breath-by-breath basis and later averaged over 10s sliding intervals for further analysis. The initial power output was 50W for 4min, followed by increases of 25W each 2min until 80% of maximal HR predicted by age was reached, then 25W increments each 1min until volitional exhaustion. The criteria for V˙O2max were RER > 1.1, plateau in V˙O2 (change of <100ml/min in the last three consecutive 20-s averages), and a HR within 10 beats/min of the maximal level predicted by age. Results of this test (see **SI Fig4** for VO2max distribution) were used to select power output for subsequent constant tests, based on the relationship between V’O2 and power output.

Blood samples: Overall 12 ml of blood were collected before and after the rest or exercise periods. Seven ml of blood were collected into a BD Vacutainer clot-activator tube (CAT), allowed to clot for 30 minutes at room temperature and centrifuged at 1100g for 15 minutes at 4°C. Serum was collected from the supernatant in aliquots of 200μl and frozen at -80°C until analysis. The other 5ml of blood were collected into a BD Vacutainer K_2_EDTA 5.4mg tube and centrifuged immediately at 8009g for 10 min. Plasma was collected from the supernatant in aliquots of 200μl frozen at -80°C until analysis. All samples were centrifuged in a Heraeus Biofuge Stratos (ThermoFisher) centrifuge. The Quantikine ELISA Human Free BDNF kits (R&D sytems) were used to quantify serum BDNF via an enzyme-linked immunosorbent assay (ELISA) following the manufacturer’s instructions.

AEA and 2-AG were extracted from 100μl of plasma by liquid-liquid extraction, and then separated by liquid chromatography (Ultimate 3000RS, Dionex, CA, USA). Analyses were performed on a 5500 QTrap® triple quadrupole/linear ion trap (QqQLIT) mass spectrometer equipped with a TurboIon-SprayTM interface (AB Sciex, Concord, ON, Canada) as described previously (20, 21).

#### Functional MRI analysis

Conventional MRI analysis: Functional images were analyzed using SPM12 (Wellcome Department of Imaging Neuroscience, London, UK). This analysis included standard preprocessing procedures: realignment, slice timing to correct for differences in slice acquisition time, normalization (images were normalized to an MNI template), and smoothing (with an isotropic 8-mm FWHM Gaussian kernel) – except for the decoding analysis where we used unsmoothed images (see below). While scanning was not performed right after physical exercise or rest, but about 1h later, we nevertheless performed corrections to regress out potential physiological artifacts from heart rate and breathing using Retroicor (14) and RVHcorr (15, 16), respectively.

Psychophysiological Interaction analysis: Psychophysiological interaction (PPI) analysis was computed to test the hypothesis that functional connectivity between the hippocampal seed region identified in the test session (at 18, -38, -8) and the rest of the brain differed during the retest session as a function of Exercising Condition in which a given trial was initially presented. Therefore, we took as psychological factor the contrast between Moderate intensity exercise and Rest, irrespective of trial type (direct, inference 1 and inference 2 trials). A new linear model was prepared for PPI analyses at the individual level, using three regressors. The first regressor represented the psychological factor, composed of moderate intensity exercise vs rest hits. The second regressor was the activity in the hippocampal seed region. The third regressor represented the interaction of interest between the first (psychological) and the second (physiological) regressor. To build this regressor, the underlying neuronal activity was first estimated by a parametric empirical Bayes formulation, combined with the psychological factor and subsequently convolved with the hemodynamic response function (22). The model also included movement parameters as regressors of no interest. A significant psychophysiological interaction indicated a change in the regression coefficients between any reported brain area and the reference region, related to the correct retrieval after moderate intensity exercise versus after rest trials. Next, individual summary statistic images obtained at the first-level (fixed-effects) analysis were spatially smoothed (6mm FWHM Gaussian kernel) and entered a second-level (random-effects) analysis using ANOVAs to compare the functional connectivity between groups.

Decoding analysis: A decoding procedure was performed on unsmoothed data. For each session of each participant, we extracted the timeseries of all voxels within the bilateral hippocampus region of interest, which was defined in the Anatomy toolbox as the union of CA1, CA2, CA3 and DG regions. Timeseries were detrended and demeaned, then the movement parameters obtained from realignment and breathing parameters from Retroicor and RVHcorr were regressed out. Mimicking the analysis of Van Dongen et al. (23), we included correct, incorrect and control trials in the analysis. Estimates of the BOLD response for each single trial were then computed to obtain a “voxel by trial matrix”, from which the mean BOLD response for each type of trial (correct, incorrect, and control) was computed per voxel. Decoding accuracy was obtained by first applying a leave one out procedure, computing a mean “voxel by trial type matrix” for all participants but one. A standard cross-validation procedure was then performed for each trial of the left out participant and it was classified as a trial type where the highest Pearson R correlation was found. Overall, we obtained percentages of trials classified as correct trials, incorrect trials, and control trials for each trial type, which were used for statistical analysis of sensitivity (true positive rate) and specificity (true negative rate). To assess whether there was a laterality effect in the hippocampus, we subsequently ran analyses using the left and right hippocampus as separate regions of interest (see **SI Fig3**).

### Data and code availability

The data that supports these findings and the custom code used in this study are available from the corresponding author upon reasonable request.

### SI Figures

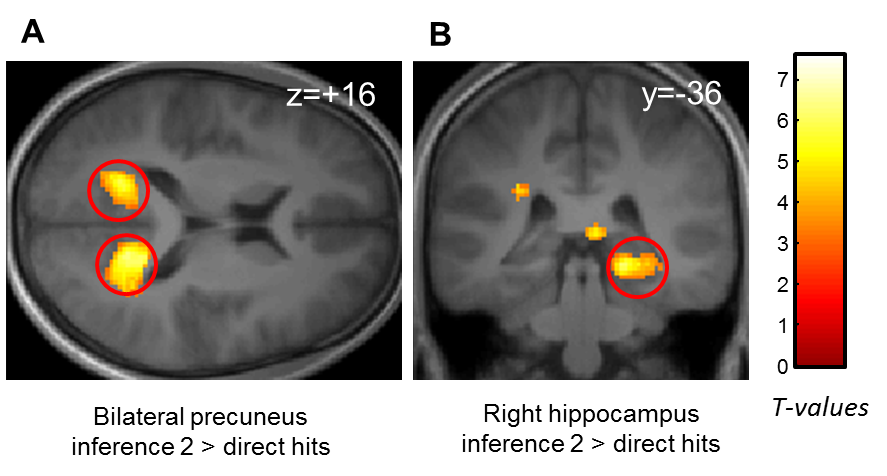

#### Fig. S1 - Brain correlates of increasing Relational Distance

**A)** Bilateral precuneus activation for increasing Relational Distance (inference 2 hits > direct hits). **B)** Right hippocampal activation for increasing Relational Distance (inference 2 hits > direct hits) [z score=4.35 (18, -38, -8), p<0.05 SVC].

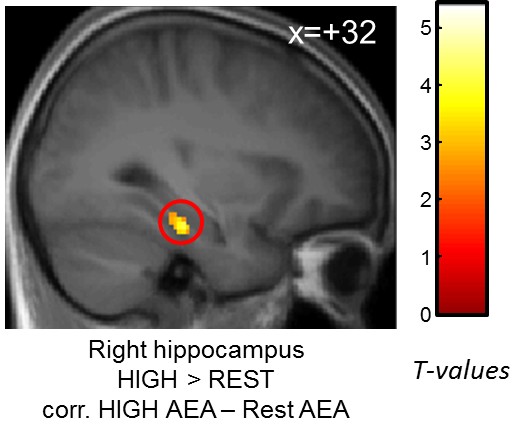

#### Fig. S2 - Hippocampal response correlated with endocannabinoid increase after high intensity exercise

Increased right parahippocampal (extending into hippocampus) response [z-score=4.08 (32, -24, -18), p<0.05 SVC] for hits after high intensity exercise compared to hits after rest correlated with the increase in AEA level after high intensity exercise.

Activation map displayed on the mean T1 anatomical scan of the whole population.

For display purposes, hippocampal activation is thresholded at p<0.005.

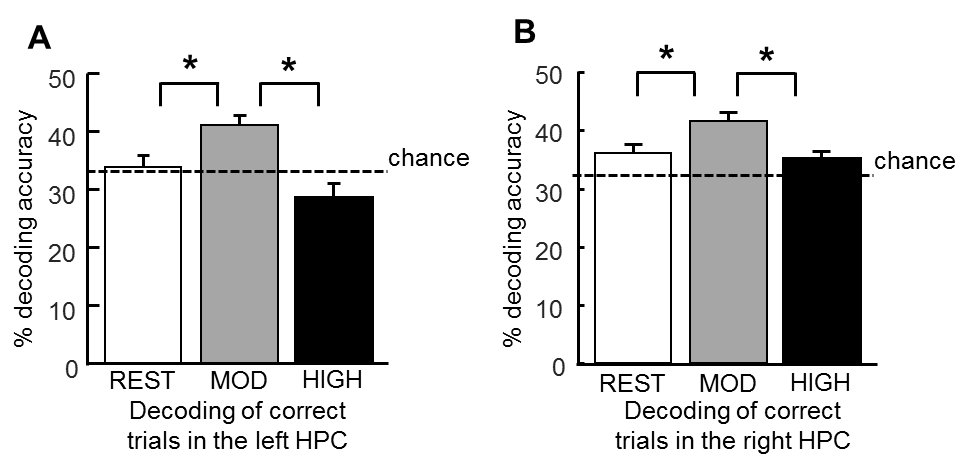

#### Fig S3 – Better decoding accuracy in both left and right hippocampi after moderate intensity exercise

**A)** Better sensitivity of decoding of correct trials in the left hippocampus after moderate exercise than rest and high intensity exercise. ANOVA F(2, 34)=7.60, p=0.002, post-hoc p_mod-rest_=0.03 p_mod-high_=0.001. Decoding after both rest and high intensity exercise is not different from chance level (both p>0.05), while being above chance level after moderate intensity exercise (t(17)=4.38, p<0.001). B) Better sensitivity of decoding of correct trials in the right hippocampus after moderate exercise than rest and high intensity exercise. ANOVA F(2, 34)=5.24, p=0.01, post-hoc p_mod-rest_=0.01 p_mod-high_=0.01. Decoding after both rest and high intensity exercise is not different from chance level (both p>0.05), while it is above chance level after moderate intensity exercise (t(17)=5.15, p<0.001).

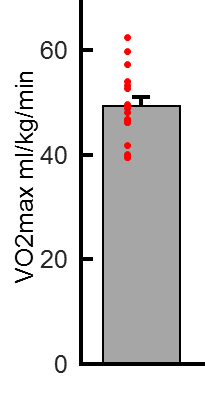

#### Fig. S4 – Distribution of VO2max measures

| **Brain Region** | **Lat.** | **cluster size** | **unc. p-value** | **SVC p-value¤** | **peak T** | **peak Z** | **X** | **Y** | **Z** |
| --- | --- | --- | --- | --- | --- | --- | --- | --- | --- |
| **Increasing relational distance (inference 2 hits > direct hits)** | | | | | | | | | |
| Precuneus | Right | 579 | 7.9E-07 |  | 7.16 | 4.80 | 16 | -46 | 14 |
| Precuneus | Left | 366 | 8.4E-06 |  | 5.92 | 4.30 | -18 | -54 | -10 |
| **Hippocampus** | **Right** | **190** | **4.2E-06** | **0.02** | **6.27** | **4.35** | **18** | **-38** | **-8** |
| Subiculum | Right | 15 | 3.7E-04 | 0.01 | 4.10 | 3.37 | 26 | -28 | -20 |
| Lingual gyrus | Right | 50 | 3.6E-05 |  | 5.20 | 3.97 | 16 | -82 | -6 |
| Occipital gyrus | Right | 13 | 4.9E-04 |  | 3.97 | 3.29 | -12 | -92 | 0 |
| **Moderate intensity exercise > rest corr. with changes in AEA** | | | | | | | | | |
| **Hippocampus** | **Right** | **13** | **1E-04** | **0,015** | **5.11** | **3.72** | **38** | **-14** | **-20** |
| **High intensity exercise > rest corr. with changes in AEA** | | | | | | | | | |
| Parahippocampus | Left | 35 | 9.4E-6 |  | 6.54 | 4.28 | -30 | -28 | -18 |
| Parahippocampus | Right | 29 | 2.7E-5 |  | 5.99 | 4.08 | 34 | -24 | -22 |
| **Hippocampus (extending from parahippocampus above)** | **Right** | 29 | 2.7E-5 | **0,02** | 4.95 |  | **32** | **-24** | **-18** |
| Middle Occipital Gyrus | Right | 21 | 6.5E-05 |  | 5.36 | 3.83 | 42 | -78 | 36 |

#### Table S1 – Activated brain regions at test

**¤ Activations in the hippocampal formation corrected using an anatomical mask (see Methods)**
